# Scaling of gene transcriptional gradients with brain size across mouse development

**DOI:** 10.1101/2020.06.04.135525

**Authors:** Lau Hoi Yan Gladys, Alex Fornito, Ben D. Fulcher

## Abstract

The structure of the adult brain is the result of complex physical mechanisms acting through development. These physical processes, acting in threedimensional space, mean that the brain’s spatial embedding plays a key role in its organization, including the gradient-like patterning of gene expression that encodes the molecular underpinning of functional specialization. However, we do not yet understand how the dramatic changes in brain shape and size that occur in early development influence the brain’s transcriptional architecture. Here we investigate the spatial embedding of transcriptional patterns of over 1800 genes across seven time points through mousebrain development using data from the Allen Developing Mouse Brain Atlas. We find that transcriptional similarity decreases exponentially with separation distance across all developmental time points, with a correlation length scale that follows a powerlaw scaling relationship with a linear dimension of brain size. This scaling suggests that the mouse brain achieves a characteristic balance between local molecular similarity (homogeneous gene expression within a specialized brain area) and longer-range diversity (between functionally specialized brain areas) throughout its development. Extrapolating this mouse developmental scaling relationship to the human cortex yields a prediction consistent with the value measured from microarray data. We introduce a simple model of brain growth as spatially autocorrelated gene-expression gradients that expand through development, which captures key features of the mouse developmental data. Complementing the well-known exponential distance rule for structural connectivity, our findings characterize an analogous exponential distance rule for transcriptional gradients that scales across mouse brain development, providing new understanding of spatial constraints on the brain’s molecular patterning.

## Introduction

Brain structure is the result of physical mechanisms playing out through development, shaped by genetic and environmental factors. In particular, the embedding of neural tissue in physical space places major constraints on neural organization. For example, the connection probability between pairs of neural elements decays with their separation distance in *C. elegans* (1), mouse (2, 3), rat (4), zebrafish (5), non-human primate (6), and human (7), and is frequently characterized as an exponential distance rule (8–10). Indeed, very simple spatial rules can explain much of the high-order statistics of connectome topology, including its modularity, longtailed degree distribution, local network motif probabilities, and the existence of a network core (7, 9, 11–15). Spatial proximity also plays a strong role in shaping gradients of gene expression, with nearby areas exhibiting more similar gene-expression profiles than more distant areas in the head neurons of *C. elegans* (1), the mouse brain (2, 16), and the human cortex (17–22).

The spatial embedding of gene-expression patterns provides important clues about how the brain’s functional specialization is organized. Macroscopic functional gradients across the human cortex maximize the spatial distance between areas functionally involved in sensory perception and those involved in integrative cognition (23, 24). These hierarchical functional gradients are underpinned by a corresponding variation in structural microarchitecture (25–27). But given the constraint of a fixed brain size, how does the brain strike the right balance between local molecular similarity (similar geneexpression patterns supporting regional specialization of function), with inter-regional molecular diversity, which supports functional heterogeneity. Furthermore, how does the spatial embedding of transcriptional gradients emerge through development? Do different spatial constraints dominate early in development compared to later? Although macroscopic brain organization in adult is often attributed to gradients set up during development (e.g., in transcriptional factors) (28), to our knowledge, no study has analyzed the spatial embedding of gene expression through development.

The Allen Developing Mouse Brain Atlas (ADMBA) (29) is a comprehensive database of gene-expression across mouse-brain development. The atlas contains *in situ* hybridization measurements for approximately 2100 genes at seven developmental time points, available in a three-dimensional reference space registered to an anatomical reference atlas at each developmental time point. Many analyses of gene transcription through mouse-brain development have shed light on how the spatiotemporal variation of specific genes through development shape different aspects of brain structure (29–32). But these developmental atlas data have not previously been analyzed from the viewpoint of spatially embedded gradients of transcriptional similarity. By characterizing gradients, we can address global questions about transcriptional patterning through development in a parcellation-free way that does not involve the difficult task of tracking anatomically defined areas.

Here we study how correlated gene expression (CGE), defined as the similarity in the gene-expression signatures of two points in the brain, is spatially embedded across mouse-brain development. Matching the well-known exponential distance rule for connectivity, we find an exponential decay in CGE at each developmental time point, with a spatial correlation length that scales with a linear dimension of brain size. Extrapolating the relationship fitted on developing mouse-brain data allows us to obtain an estimate consistent with the measured correlation length of CGE in the human cortex. Using a simple model, we show that the patterns are consistent with brain growth in which spatially autocorrelated gene-expression gradients expand with brain size over time.

## Methods

### Data

We analyze data from the ADMBA, which contains *in situ* hybridization (ISH) expression estimates for approximately 2100 genes in C57Bl/6J mice across seven developmental stages, spanning both embryonic (E11.5, E13.5, E15.5, and E18.5) and postnatal (P4, P14, and P28) stages (29). These genes were selected to encompass relevant transcription factors, neurotransmitters and their receptors, cell-type or regional marker genes, genes relevant to developmental signaling pathways, and genes corresponding to common drug targets or implicated in neurodevelopmental disorders (29). The raw imaging data, encompassing 434 946 brain sections of high-resolution ISH, are registered to a three-dimensional mouse-brain atlas at each time point. We accessed this registered data at the voxel level via the Allen Software Development Kit (SDK, 2015). Our code for retrieving data (in python) and for data processing and analysis (Matlab) is available at https://github.com/NeuralSystemsAndSignals/DevelopingMouse. At each time point, we retrieved expression data for all available genes. We used sagittal section data for its comprehensiveness, which had coverage across the left hemisphere at each time point. Expression data for each gene was obtained as ‘expression energy’ (ISH signal intensity) in each voxel in the threedimensional grid. Distances between pairs of voxels were computed as Euclidean distances in three-dimensional space using the spatial resolution of the coordinate grid at each time point: E11.5 (0.08mm), E13.5 (0.1mm); E15.5 (0.12mm), E18.5 (0.14mm), P4 (0.16mm), P14 (0.2mm), and P28 (0.2mm).

### Filtering and normalization

At each developmental time point, we first filtered out voxels that were labeled as spinal cord or were unannotated, and labeled voxels according to four anatomical masks: ‘forebrain’, ‘midbrain’, ‘hindbrain’, ‘dorsal pallium’. At each time point, we retained genes with valid expression (i.e., not a missing value) for at least 70% of voxels, resulting in different numbers of good-quality genes across time: E11.5 (2038 genes), E13pt5 (2085), E15pt5 (1861), E18pt5 (2053), P4 (2056), P14 (2015), and P28 (2063). These small differences in the number of genes across time did not affect our results: we obtained similar findings when restricting our analysis to a consistent subset of 1861 genes that had good-quality data across all time points. We then retained only voxels with valid expression data for at least 70% of these genes. The number of voxels varied across time as 5034 (E11.5), 9473 (E13.5), 11 314 (E15.5), 11313 (E18.5), 19754 (P4), 21579 (P14), and 24 822 (P28). We generated a voxel × gene expression matrix containing voxels and genes that survived these filtering steps.

Raw expression values measured using *in situ* hybridization are not comparable between genes (33). We accounted for this at each time point by normalizing expression values of each gene using an outlier-robust sigmoidal transformation, *S*(*x*) = [1+exp(−(*x* – *x*_med_)/1.35*x*_iqr_)]^−1^, where *x*_med_ is the median of the expression vector, *x*, for a given gene, and *x*_iqr_ is its interquartile range (17, 34). For visualization, this normalization was followed by linear rescaling to the unit interval. The resulting normalized voxel × gene matrix, *G_t_*, at a given time point, *t*, was used for further analysis.

### Quantifying the spatial embedding of CGE

To understand the spatial embedding of gene-expression patterns, we analyzed the distance-dependence of correlated gene expression (CGE) at each developmental time point. CGE is a measure of gene-expression similarity between a pair of voxels and was computed as the Pearson correlation coefficient between normalized gene-expression vectors (rows of a given voxel × gene matrix, *G_t_*, defined above).

At each time point, we computed CGE for every pair of voxels. Because the distribution of pairwise distances is concentrated near lower distances, we wanted to prevent the exponential fitting from being biased towards fitting these smaller length scales, but to instead fit well across the full range of distances. To do this, we first performed an equiprobable distance binning of the data using twenty bins, and computed the mean CGE value in each bin (and summarizing each bin as its center). To this binned data, we then fitted a three-parameter exponential function, CGE(*d*) = *A*exp(−*d*/λ) + *f*_0_, using nonlinear least squares and identified the fitted parameters as: spatial correlation length, λ, strength *A*, and offset, *f*_0_. Adjusted *R*^2^ was used as a goodness-of-fit statistic. Note that for self-correlation, at *d* = 0, CGE = 1, and thus fitting data that includes self-correlations should have a fixed strength parameter *A* =1. Here we excluded self-correlations and allowed the strength parameter, *A*, to vary. When characterizing distances relative to brain size, as *d*_rel_ = *d*/*d*_max_, we defined *d*_max_ as the maximum extent along the anterior–posterior axis of the brain. For computational efficiency, we restricted our analysis to a random subset of 1000 voxels at each developmental time point. Our results do not strongly depend on this spatial downsampling (e.g., λ estimates stabilized for samples of approximately 500 voxels, cf. Fig. S4).

Note that our definition of λ as a length scale (m) is consistent with its conventional use to denote wavelength in physics (units of length), but differs by an inverse relationship from its use as a decay rate with units of inverse length (m^−1^) used to characterize the exponential distance rule for structural connectivity (4, 8–10). To estimate and visualize the dominant expression pattern across all genes in matrices, *G_t_*, we used principal components analysis, accounting for missing values using alternating least squares.

### Modeling

To better understand patterns found in the empirical data, we simulated simple spatial models of gene expression patterning and brain growth. At each developmental time point, we first simulated geneexpression patterns for each of the 1861 genes. We first defined the geometry by approximating the brain as a rectangular prism with dimensions (anterior–posterior × superior–inferior × left–right) equal to that of the developmental reference atlas at that time point: 3.52 × 4.72 × 1.20mm (E11.5), 5.70 × 3.30 × 1.20mm (E13.5), 6.60 × 3.72 × 1.68mm (E15.5), 7.84 × 4.06 × 2.10mm (E18.5), 10.56 × 5.44 × 3.04mm (P4), 12.80 × 6.60 × 4.20 mm (P14), and 13.60 × 7.40 × 4.20 mm (P28). We formed a grid over this idealized geometry by dividing each dimension into 50 equally spaced points. We then generated a random spatially autocorrelated map independently for each of 1861 genes using a spatial-lag model (25, 35, 36). The model defines a dependence between samples using an exponential kernel as *W_ij_* = exp(−*d_ij_*/*d*_0_), for pairwise distances, *d_ij_*, and a characteristic spatial scale, *d*_0_. From this weighting matrix, *W_ij_*, an autocorrelated spatial map can be generated as *x_i_* = (*I* + *ρW_ij_*)*u_j_*, where *I* is the identity and 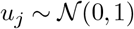 is i.i.d. Gaussian noise. This defines the second model parameter: the spatial autocorrelation strength, *ρ*. By setting *d*_0_ as a fixed fraction of brain size, 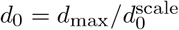, we simulated a brain-size scaling rule consistent with a linear rescaling of space (spatial expansion through development). Values for *ρ* and 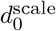 were set through a gradient descent optimization procedure to best fit the empirical values of λ and *A*, yielding *ρ* = 0.416 and 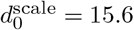. Some illustrative examples of the expression patterns generated are plotted in Fig. 6A.

Note that at small distances, all grid points are included in the CGE calculation. But due to the grid’s finite size, beyond a critical distance, only a subset of grid points are included in the CGE calculation (e.g., in the extreme case of the maximum distance, only the corners of the cube are included). This finite-size spatial sampling effect can bias the CGE(*d*) curve at large distances. For example, distance bins that are affected by this are labeled pink in Fig. 6B.

For each simulated gene-expression dataset, we performed the same processing and analysis methodology as was applied to the empirical data: (i) a random subsample of 1000 points was taken; (ii) voxel × gene expression data were normalized using a scaled sigmoid; (iii) CGE was computed; (iv) CGE(*d*) data were binned using 20 equiprobable bins; (v) exponential decay was fit. We repeated the process 50 times to estimate error bars on each parameter estimate under the sources of stochasticity in the model: the generation of spatially autocorrelated gene-expression maps, and the random spatial subsampling. We were then able to compare the parameter estimates from this model with the empirical data. Code for the modeling is available at https://github.com/NeuralSystemsAndSignals/DevelopmentalExpressionModeling.

## Results

Our approach to investigating the spatial embedding of transcriptional gradients across mouse-brain development is shown schematically in Fig. 1. At each of the seven time points in the ADMBA (29), shown in Fig. 1A, we obtained the expression of 1861 genes across voxels of the mouse brain (Fig. 1B). We then computed the correlated gene expression (CGE) for each pair of voxels as the Pearson correlation coefficient between their expression signatures (rows of the matrix in Fig. 1B). We fitted an exponential function to the decay of CGE(*d*) as a function of separation distance, *d*, shown in Fig. 1C, to quantify how transcriptional similarity is spatially embedded. Variation in the parameters of this fit, particularly that of the characteristic length scale over which genes are correlated, λ, tell us how the properties of gene-expression gradients vary across mouse-brain development.

**Fig. 1.**
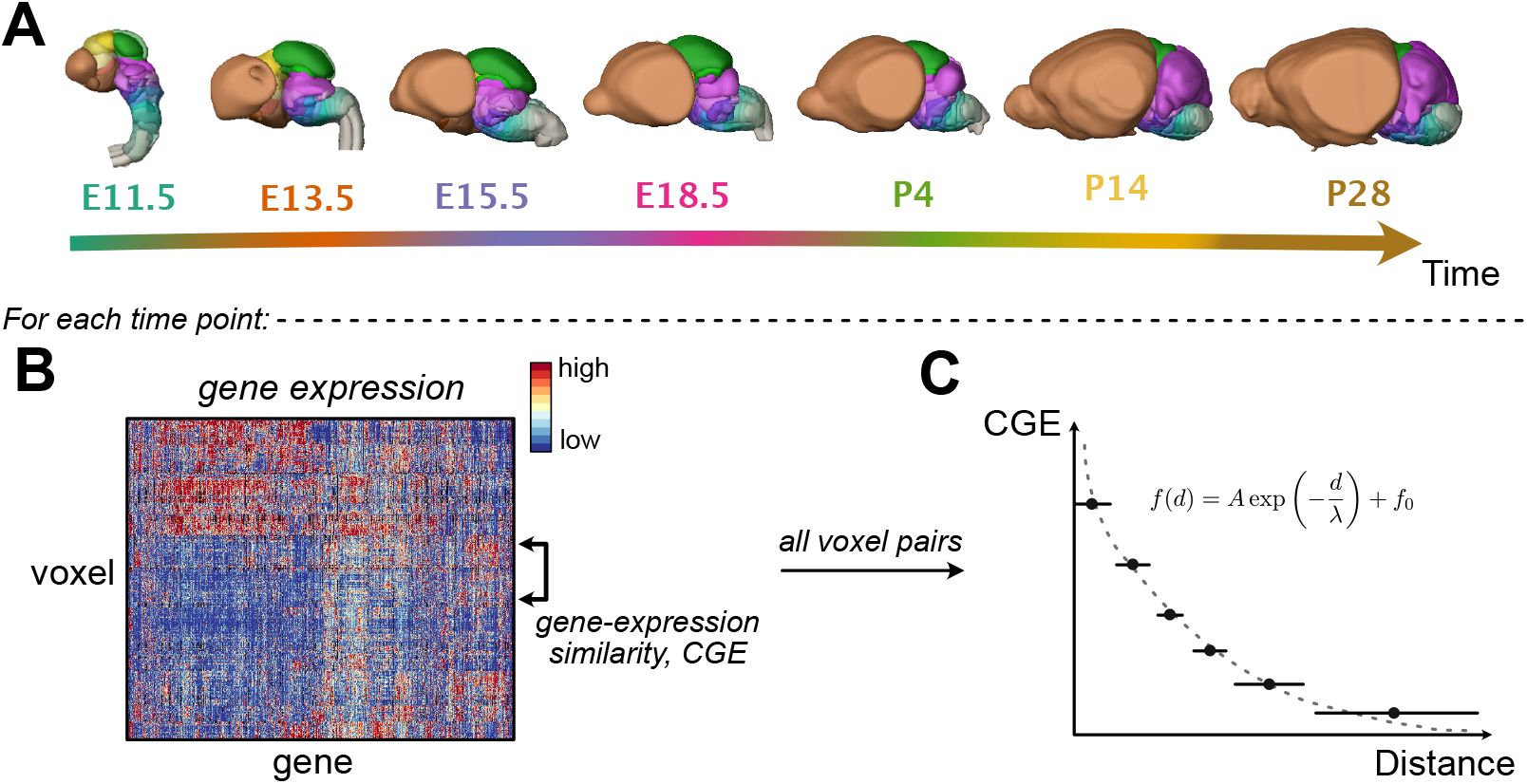
Schematic analysis pipeline. **A** We analyze geneexpression atlases across seven developmental time points. **B** At each time point, we created a voxel × gene expression matrix (data shown for E11.5). We then compared the correlated gene expression (CGE) between each pair of voxels as a function of their physical separation distance, *d*. **C** Plotting CGE(*d*) across equiprobable distance bins allowed us to analyze the spatial embedding of gene expression at each time point. We fitted an exponential form CGE(*d*) = *A* exp(−*d*/λ) + *f*_0_. Here we focus on the correlation length, λ, and how it scales with brain development.

### Low-dimensional spatial gradients

To better understand the dominant patterns of gene expression in the mouse brain, we first visualized voxel × gene expression matrices for each time point, and projected their leading principal components into brain space. Examples for two selected time points are shown in Fig. 2 (see Fig. S1 for all time points). Gene-expression matrices have been reordered to put genes with similar expression close to each other (columns of the expression matrices in Figs 2A,C) and, within three main divisions—forebrain, midbrain, and hindbrain—voxels with similar expression signatures close to each other. Visually, we find interesting gene-expression patterns that clearly differ between the three major brain divisions. The similarity-based ordering of columns in these expression matrices makes the low-dimensionality of the expression data visually clearer: we see how large numbers of genes display highly correlated expression patterns. The leading principal component of the expression data (computed using alternating least squares to account for a small proportion of missing data) is annotated to the left of Figs 2A,C, and visually follows the types of dominant gene-expression patterns that capture the most variance across all genes.

**Fig. 2.**
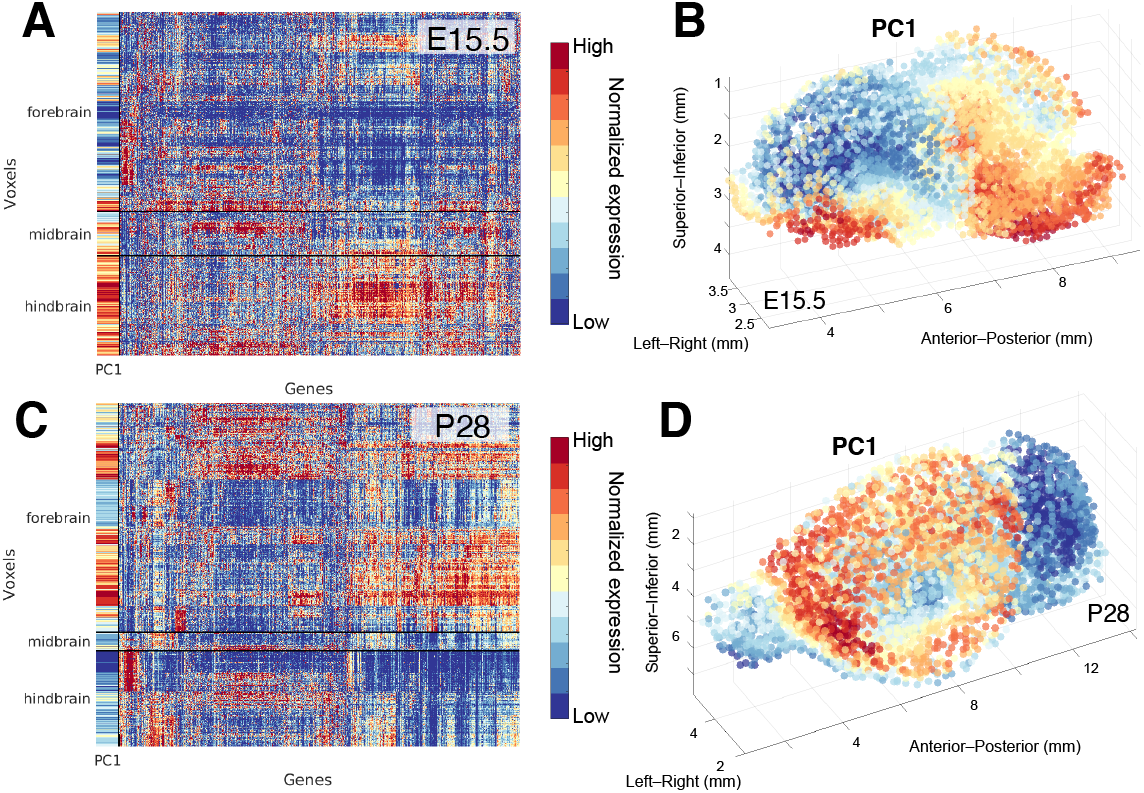
Developmental mouse gene-expression patterns display low-dimensional spatial gradients. Here we plot results for E15.5 and P28 (see Fig. S1 for all time points). We show voxel × gene expression matrices as heat maps for E15.5 and P28 in **A** and **C**, respectively. All brain voxels and genes passing quality-control criteria are shown (11314 × 1851 for E15.5, and 24797 × 1846 for P28). Each gene has been normalized according to an outlier-robust sigmoidal transform, followed by rescaling to the unit interval, from relative low expression for a given gene (blue) to relative high values (red). To visualize the structure in these data, genes have been reordered by their correlation similarity, and within each labeled subdivision (forebrain, midbrain, hindbrain), voxels have been similarly reordered. Horizontal black lines distinguish voxels from these three major subdivisions. The leading principal component of each matrix is labeled as ‘PC1’, and visually follows the dominant expression pattern across genes. These leading principal components are plotted in three-dimensional brain space in **B** and **D** for E15.5 and P28, respectively. To aid visualization of the three-dimensional structure, a random subset of 5000 voxels is plotted with transparency. There is clear spatial autocorrelation in these low-dimensional expression gradients in the developing mouse brain, which we quantify in this work.

To determine whether these low-dimensional components are spatially embedded as gradients, we plotted them in Figs 2B,D for E15.5 and P28, respectively. We find clear spatial autocorrelation in these lowdimensional spatial gradients, an effect that is also seen across the other developmental time points (Fig. S1), and for lower-order principal components (not shown). Having built some intuition for the existence of lowdimensional spatial gradients of gene-expression in the developing mouse brain, in the remainder of this study we aim to quantitatively these gradients across development using CGE(*d*) curves.

### Distance dependence of correlated gene expression

Binned CGE(*d*) data with exponential fits are shown for each of the seven developmental time points in Figs 3A–G. The three-parameter exponential form captured the data well at all time points (all adjusted *R*^2^ > 0.96). This exponential distance-dependence of gene-expression similarity provides a quantitative way to characterize the gradient-like expression patterns shown in Figs 2B,D.

**Fig. 3.**
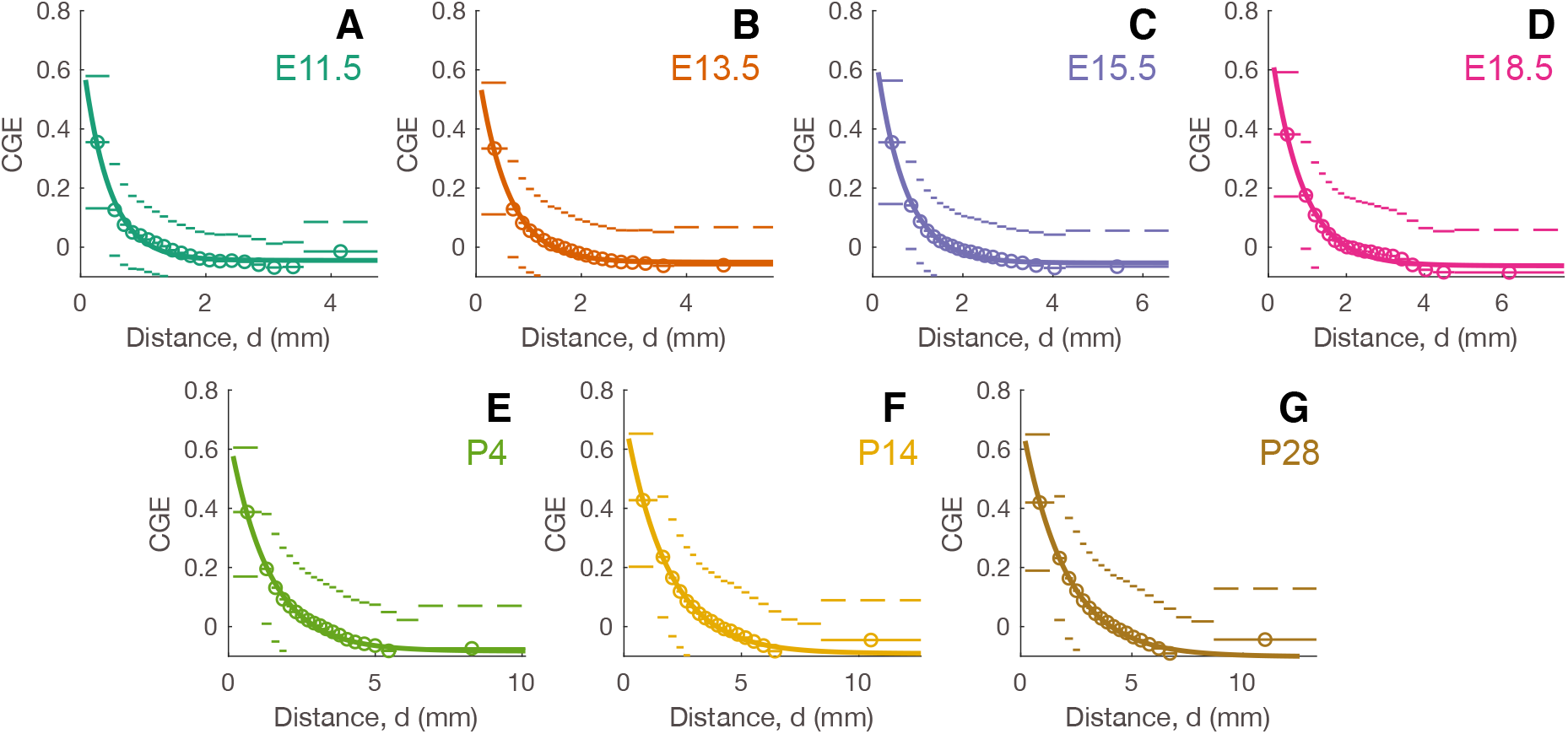
Correlated gene expression (CGE) exhibits an approximately exponential decay with distance at all developmental time points. **A–G**: Across twenty equiprobable distance bins, mean CGE of all pairs of voxels in each bin is plotted as circles, with horizontal lines showing the extent of each bin. Dashed lines indicate one standard deviation either side of the mean of each bin. Fitted exponential curves, CGE(*d*) = *A*exp(−*d*/λ) + *f*_0_, are shown solid.

We next investigated whether the CGE(*d*) curves shown in Fig. 3, computed using all voxels, vary as a function of anatomical division, focusing on the forebrain, midbrain, and hindbrain. We recomputed CGE(*d*) curves for each type of voxel pair: forebrain–forebrain, forebrain– midbrain, midbrain–midbrain, midbrain–hindbrain, or hindbrain–hindbrain, shown in Fig. S2. Most of these anatomically distinguished CGE(*d*) curves decayed with distance, but with different length scales and strengths. There were some deviations, such as an increase in midbrain–midbrain CGE at ~ 3 mm at E18.5, which causes a slight ‘hump-like’ deviation from an exponential form for the bulk CGE(*d*) curve (Fig. 3D). Consistent with an increased gene-expression similarity within each anatomical division, within-division CGE was higher than between-division CGE at a given separation distance, *d*, across all developmental time points. Indeed, within-division CGE was almost always positive, even at the longest separation distances, whereas between-division CGE often became negative (i.e., geneexpression signatures are anti-correlated), even at intermediate separation distances. This between-division anti-correlation contributes to the overall negative baseline, *f*_0_, of the exponential fits to bulk CGE(*d*) in Fig. 3. The increase in CGE towards ~ 0 at the longest separation distance bin—seen for the postnatal times P4, P14, and P28—can mostly be attributed to a decrease in anti-correlation of expression profiles between forebrain and hindbrain voxels. Postnatal time points were also distinguished by high expression similarity within forebrain voxels, even at the longest separation distances (Figs S2E–G). While the focus of this study is to investigate CGE(*d*) curves using voxels from across the whole brain, we note that this global gradient-like spatial embedding is not isotropic and non-specific, but indeed varies with anatomy.

We next investigated whether the CGE(*d*) curves vary across subsets of neuron-enriched, astrocyte-enriched, and oligodendrocyte-enriched genes (defined as being more than 1.5-fold enriched using data from Cahoy et al. (37)). Compared to using all genes, CGE(*d*) curves and relationships between fitted CGE decay parameters with brain size, remained similar for all gene subsets, indicating that the spatial embedding of gene expression does not vary strongly with genes enriched for each of these cell types.

### Scaling of transcriptional correlation length with brain size

When plotting the fitted CGE(*d*) exponential curves together (Fig. S3A) and rescaling distances by brain size, *d*_max_ (maximal anterior–posterior distance), as 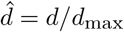 (Fig. S3B), we found that CGE(*d*) approximately collapsed onto a common relative correlation length. To quantify this relationship, we plotted the fitted transcriptional correlation length, λ, as a function of brain size, *d*_max_, in Fig. 4A. We find that λ increases with *d*_max_ through development (Pearson’s *r* = 0.98, *p* = 6 × 10^−5^). Having only seven points over less than a decade of brain sizes, *d*_max_ prevents rigorous inference of power-law scaling (38), but we have physical reasons to expect a scaling relationship across brain expansion (10). To explore this possibility, we fitted a power-law, 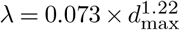, shown in Fig. 4B.

**Fig. 4.**
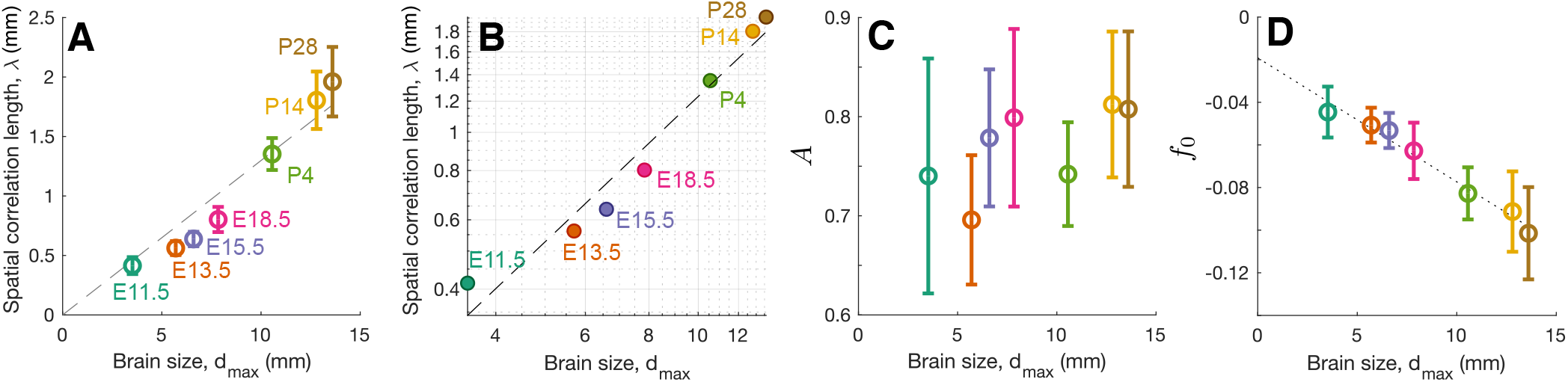
The correlation length oftranscriptional coupling, λ, increases with brain size. The three fitted parameters of the exponential form CGE(*d*) = *A* exp(−*d*/λ) + *f*_0_, vary through brain development, plotted here as a function of brain size, *d*_max_ (computed along the anterior–posterior dimension): **A** transcriptional correlation length, λ (mm) (with proportional linear fit dashed); **B** *d*_max_–λ as in A but plotted on logarithmic axes with the best power-law fit, λ = 0.0735*d*^1.22^, shown dashed; **C** spatial embedding strength, *A*; and **D** offset, *f*_0_ (with linear fit shown dotted). Each data point represents the fitted parameter value from a given time point (labeled in **A** and **B**). Error bars as symmetric 95% confidence intervals are plotted on the three linear-axis plots, A, C, and D.

The spatial embedding strength, *A*, controls the strength of the exponential relationship in CGE relative to other effects such as noise. As shown in Fig. 4C, this strength is similar across development, *A* ≈ 0.75 (Pearson’s *r* = 0.65, *p* = 0.1). The offset, *f*_0_, corresponds to the CGE in the limit of large separation distances, *d*. As plotted in Fig. 4D, *f*_0_ decreases linearly, becoming more negative across development (Pearson’s *r* = −0.99, *p* = 3 × 10^−5^). These increasingly anti-correlated geneexpression patterns at the longest separation distances are consistent with an overall increase in transcriptional diversity within the brain across development.

### Predicting the transcriptional correlation length in human cortex

We next aimed to test the power-law form for the scaling of transcriptional correlation length, λ, which suggests that the mouse brain maintains a characteristic spatial correlation length of transcriptional similarity relative to its size. We also wanted to investigate whether the relationship found across mouse-brain development could be extrapolated to make a prediction of the transcriptional correlation length, λ, in human cortex. We used existing data for human cortex, *d*_max_ ≈ 148 mm (maximal anterior–posterior distance in the Glasser et al. (39) parcellation) and 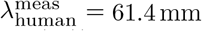 (95% confidence interval: 54.1–71.0mm (17)). The extrapolation of the power-law relationship fitted to the mouse development data, 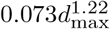, and setting *d*_max_ = 148 mm yields a predicted 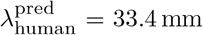 (95% CI: 12.6 mm–88.7 mm). This extrapolation is shown in Fig. 5A. The prediction is consistent with the measured value, as shown in Fig. 5B (for all voxels, labeled ‘brain’). Thus, despite differences in data type (ISH for < 2000 genes in mouse, microarray for > 20 000 genes in human), spatial discretization (at the level of voxels in mouse and a multimodal parcellation in human), we were able to use mouse developmental data to predict a key spatial property of human gene-expression gradients.

**Fig. 5.**
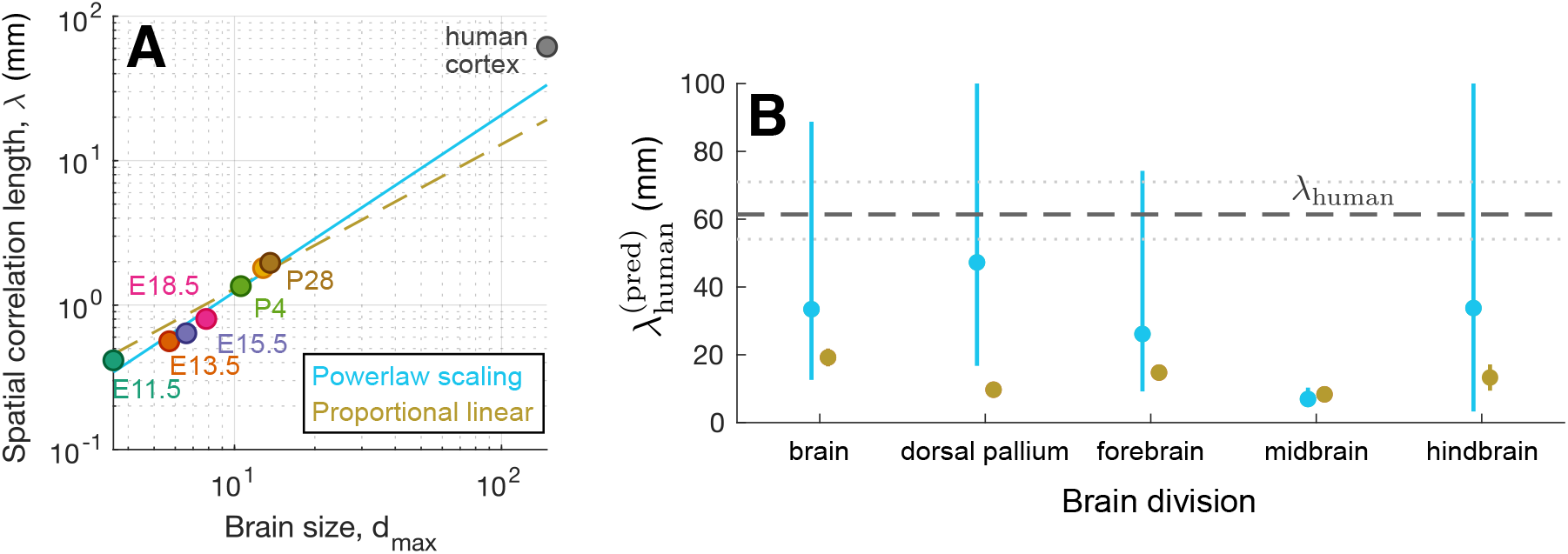
The transcriptional correlation length in human cortex, λ_human_, can be predicted by extrapolating the scaling relationship betweens λ and brain size through mouse-brain development. **A** Estimates of λ plotted as a function of brain size, *d*_max_, using all voxels at each of the mouse developmental stages. Extrapolation to the human cortex (*d*_max_ = 148 mm) is shown using a fitted power-law scaling relationship, 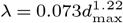 (blue), and a proportional linear relationship, λ = 0.13*d*_max_ (brown dashed). **B** We plot predictions trained from fitting to: all brain voxels, dorsal pallium (isocortex) voxels, forebrain voxels, midbrain voxels, and hindbrain voxels. We also compare the predictions of power-law scaling (blue) to those from a proportional linear scaling (brown), with 95% confidence intervals annotated.

We next investigated how this prediction varied when we restricted the fitting to a subset of voxels (dorsal pallium, forebrain, midbrain, and hindbrain), and when using a proportional linear scaling instead of the power-law form (Fig. 5B). We expected the closest estimate when using dorsal pallium voxels, as this is the structure that best corresponds to the cortex. Indeed, dorsal pallium voxels yielded the best human prediction 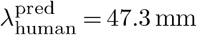 (despite the dorsal pallium not being annotated to the reference atlas at E11.5). The worst human prediction was obtained using midbrain voxels, 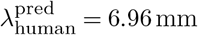. Proportional linear scaling of λ with brain size, λ ∝ *d*_max_ across mouse development consistently yielded a poorer prediction, supporting a power-law form, 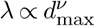, with exponent *ν* > 1. With just seven points across mousebrain development, our predictions have wide error bars; it will be important for future work to verify the results of this scaling law, and produce more constrained predictions using more comprehensive transcriptomic data, including across species.

### Modeling brain growth

To understand potential mechanisms underlying the scaling of transcriptional gradients through brain growth, we developed a simple spatial model of transcriptional patterning and tested its predictions against the empirical data characterized above. As described in Methods, we generated a random spatially autocorrelated map independently for each gene’s expression pattern, evaluated in an idealized prism geometry that matches the brain’s three-dimensional spatial extent at each developmental time point. To test whether the observed patterns are consistent with a simple rescaling of space in these expression maps, we rescaled the spatial autocorrelation length, *d*_0_, as a function of brain size, *d*_max_. The model is therefore similar to uniform, isotropic stretching of three-dimensional gene-expression patterns that exist at the earliest developmental time point (it reduces to this case when all brains have the same relative dimensions). The types of spatially autocorrelated patterns that were generated are illustrated in Fig. 6A.

**Fig. 6.**
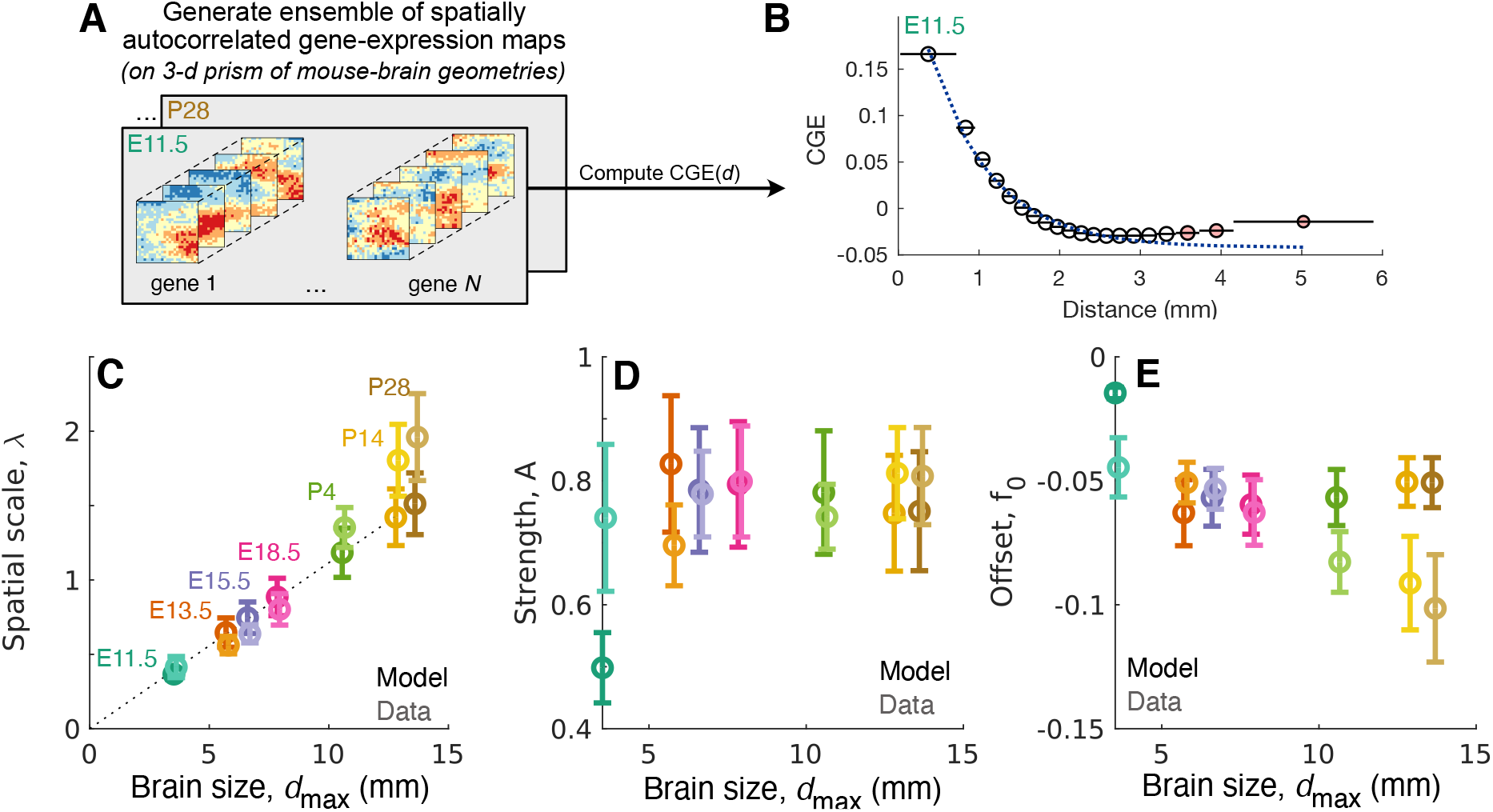
A simple spatial model for gene expression through development reproduces key features of the empirical data. **A** Illustrative ensemble of independent, spatially autocorrelated gene-expression maps evaluated on three-dimensional prisms matching the dimensions of the brain at each point in development. Examples are shown for two genes in a three-dimensional volume (expression levels shown as color). **B** Each ensemble exhibits an approximately exponential decay of correlated gene expression (CGE) with distance, *d*, as CGE(*d*) = *A*exp(−*d*/λ) + *f*_0_. Binned, model-simulated data (black) and fit (dotted blue) are plotted here for E11.5. Distance bins for which not all grid points were sampled (due to finite-size effects) are filled pink. Exponential fit parameters of the model-simulated data (shown with dark colors) and empirical data (shown with light colors) are plotted across development for: **C** spatial correlation length, λ; **D** strength, *A*; and **E** offset, *f*_0_. To aid visibility, empirical values are slightly offset horizontally.

We first verified that our model reproduces an approximately exponential dependence of CGE with distance at a given time point, shown in Fig. 6B for E11.5. Apart from a slight decrease of CGE < 0 at moderate distances, and an increase back to CGE ≈ 0 at the highest distances, we find a good fit to an exponential form. Note that the distance bins labeled pink in Fig. 6B are sufficiently large relative to brain size that they involve restricted sampling of brain space—only voxels towards the edge of the brain are able to contribute to the CGE calculation at these large distances (whereas all voxels contribute at shorter distances). The qualitative similarity between the simulated CGE(*d*) curve and the measured CGE(*d*) curves from the ADMBA data, suggest that an ensemble of genes with independent random, spatially autocorrelated transcriptional patterns can provide a good approximation for understanding how these patterns adjust through development.

After fitting the two parameters of our model, *ρ* and 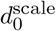, to the empirical data (see Methods), the model approximately reproduces the scaling of the correlation length, λ, with brain size, *d*_max_, seen in empirical data (Fig. 6C), and the strength, *A* (Fig. 6D).

## Discussion

Here we used data from the ADMBA to characterize how gene transcriptional gradients are spatially embedded across mouse brain development. Complementing the well-known exponential distance rules for structural connectivity in mammals (3, 4, 8, 9) that are connected by a scaling rule (10), we show that correlated gene expression is also well characterized by an exponential distance rule at each point in mouse brain development. Furthermore, the characteristic correlation length of this embedding, λ, was found to scale with a linear dimension of brain size, *d*_max_, as 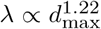 (Fig. 4). More functionally specialized brains may be expected to encode more molecularly distinctive brain areas within the constraints of its brain size, corresponding to a shorter relative λ. This scaling relationship suggests that the brain maintains a spatial trade-off between local molecular coordination (uniform gene expression within a specialized brain area) and longer-range molecular differentiation (between molecularly distinct, functionally specialized areas) across development. We were able to use this scaling law to predict the transcriptional correlation length in the adult human cortex (Fig. 5). Given that the brain-related genes exhibit strong consistency in their hierarchical patterning between mouse and human (27), it will be interesting for future work to further investigate these types of cross-species correspondences, including characterizing whether specific classes of genes drive similarities or differences in the spatial organization of gene expression across species.

A benefit of our method of characterizing CGE(*d*) curves from voxel-wise expression maps is that it does not rely on any assumptions about pre-defined anatomical parcellations or areal boundaries. Our visualizations of the voxelwise expression as clustered voxel × gene expression matrices, accompanied by their leading principal component (Figs 2A,C), show that there are low-order expression patterns that explain much of the variance across all available genes. When plotting this leading principal component in space (Figs 2B,D), we see strong gradient-like spatial autocorrelation that is quantified here (across all genes) using CGE(*d*) curves. While these analyses provide clear evidence for spatial expression gradients in the brain, they are not simple, non-specific spatial patterns, but vary markedly across anatomical divisions. This is seen most clearly when distinguishing CGE(*d*) by pairs of voxels that lie within or between each of the three major anatomical subdivisions: forebrain, midbrain, and hindbrain (Fig. S2). Consistent with characteristic expression patterns for different divisions, seen visually in Figs 2A,C, we found that within-division CGE is higher than between-division CGE at a given separation distance. Future work could take these types of analyses further to combine the benefits of a parcellation-free characterization that facilitates comparisons across development, while also investigating how properties of these spatial gradients vary as a function of anatomy. This includes the ability to score and interpret specific types of genes that contribute most to interesting patterns, such as characterizing differences between individual genes in the strength and spatial scale of their gradient-like patterning, and how these properties vary across development. The type of quantitative spatial characterization used here could also be used to construct new, parcellation-free definitions for ‘differentially expressed genes’ to test whether the hourglass pattern of transcriptional variation holds in mouse (40).

Our analysis was performed at the voxel level using *in situ* hybridization data from the ADMBA. This allowed us to explore CGE(*d*) at a much finer spatial resolution than previous analyses of correlated gene expression, performed at the level of parcellations in adult mouse brain (2) and human cortex (17). Nevertheless, each spatial element in our analysis still contains transcriptional contributions from a large number of molecularly distinct brain cells. The advent of three-dimensional *in situ* single-cell transcriptome profiling (41–44), which preserves spatial and transcriptomic information at the cellular level, will allow future analyses of correlated gene expression to be performed at even finer spatial scales. For example, this could allow testing of whether the bulk scaling rule for CGE characterized here applies at the finer scale of neuronal circuits. As transcriptomic atlas data become available for other species, such as *Drosophila melanogaster* (45, 46) and zebrafish (47), we will also be able to more comprehensively compare the spatial organization of the brain’s molecular diversity, and assess the cross-species validity of the scaling rule proposed here.

A simple model of independent, spatially autocorrelated transcriptional maps that scale with brain size (similar to the isotropic expansion of transcriptional patterns set up early in development) reproduced key features of CGE(*d*) scaling observed empirically. We kept the spatial autocorrelation scale for our simulated geneexpression maps at a fixed proportion of brain size, consistent with linear, isotropic brain growth (but allowing us to use realistic brain dimensions from empirical measurements). Our results are thus consistent with molecular specialization occurring very early in development, with brain expansion stretching the correlation length through development. While our simple isotropic model reproduces much of the CGE(*d*) relationship, our model is only evaluated against these coarse-grained properties, having blurred the contribution of the many individual genes that display precise developmental spatiotemporal programs, and the variation in these CGE(*d*) curves as a function of anatomy (Fig. S2). It will thus be important for future work to precisely investigate and model these important properties of brain development not considered here. By characterizing a generic variation in CGE, the current work may have a role to play in this research effort. For example, genes that most strongly counter the generic trends in CGE(*d*) could be considered as candidates for playing a more specific, targeted role in shaping brain development, facilitating data-driven identification of genes that play important developmental roles.

## ACKNOWLEDGEMENTS

The authors would like to thank Aurina Arnatkevičiūtė for assisting with human expression data.

## Supplementary Information

**Fig. S1.**
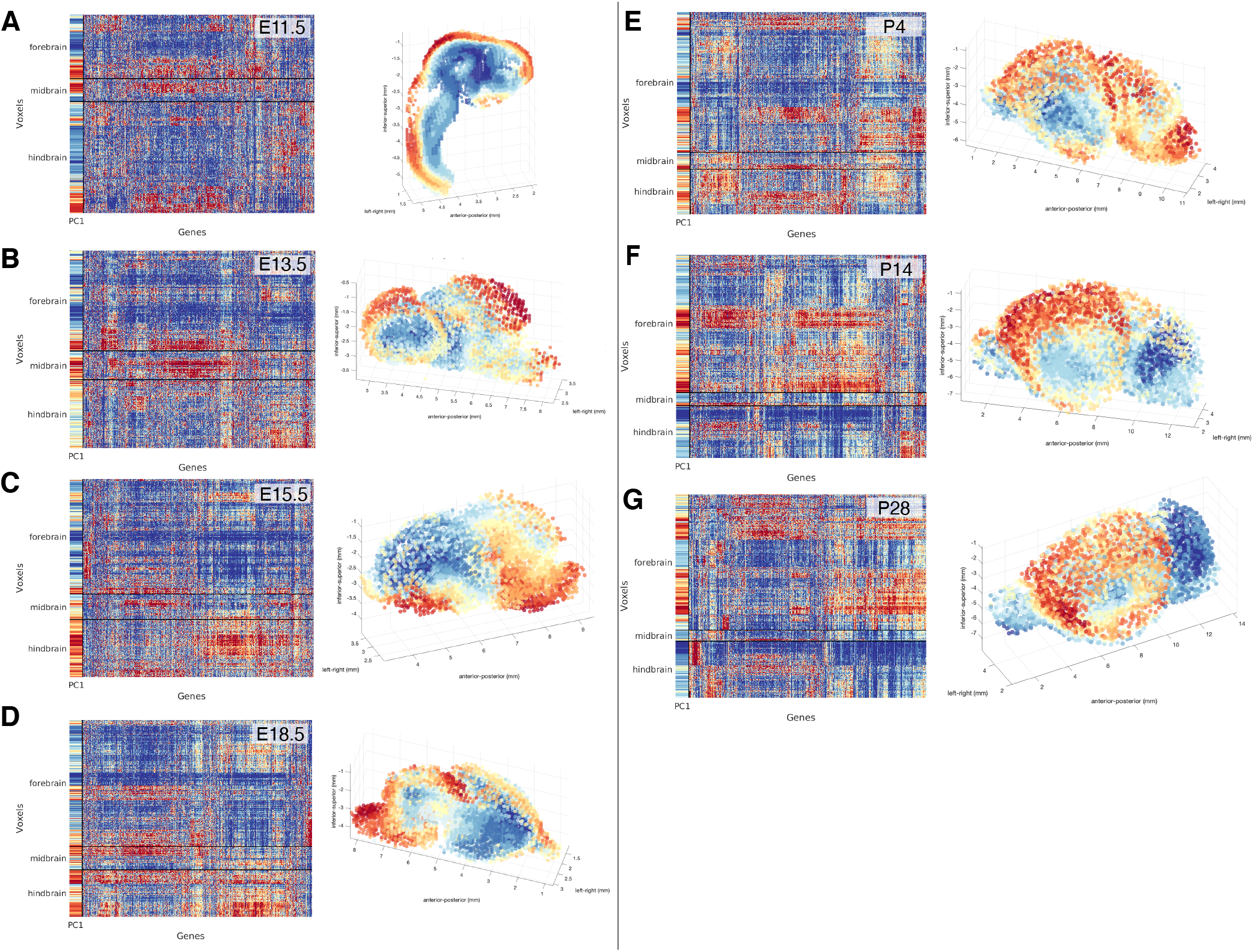
Voxel × gene expression matrices and spatial projections of their leading principal component at each developmental time point. Plots are as described in Fig. 2.

**Fig. S2.**
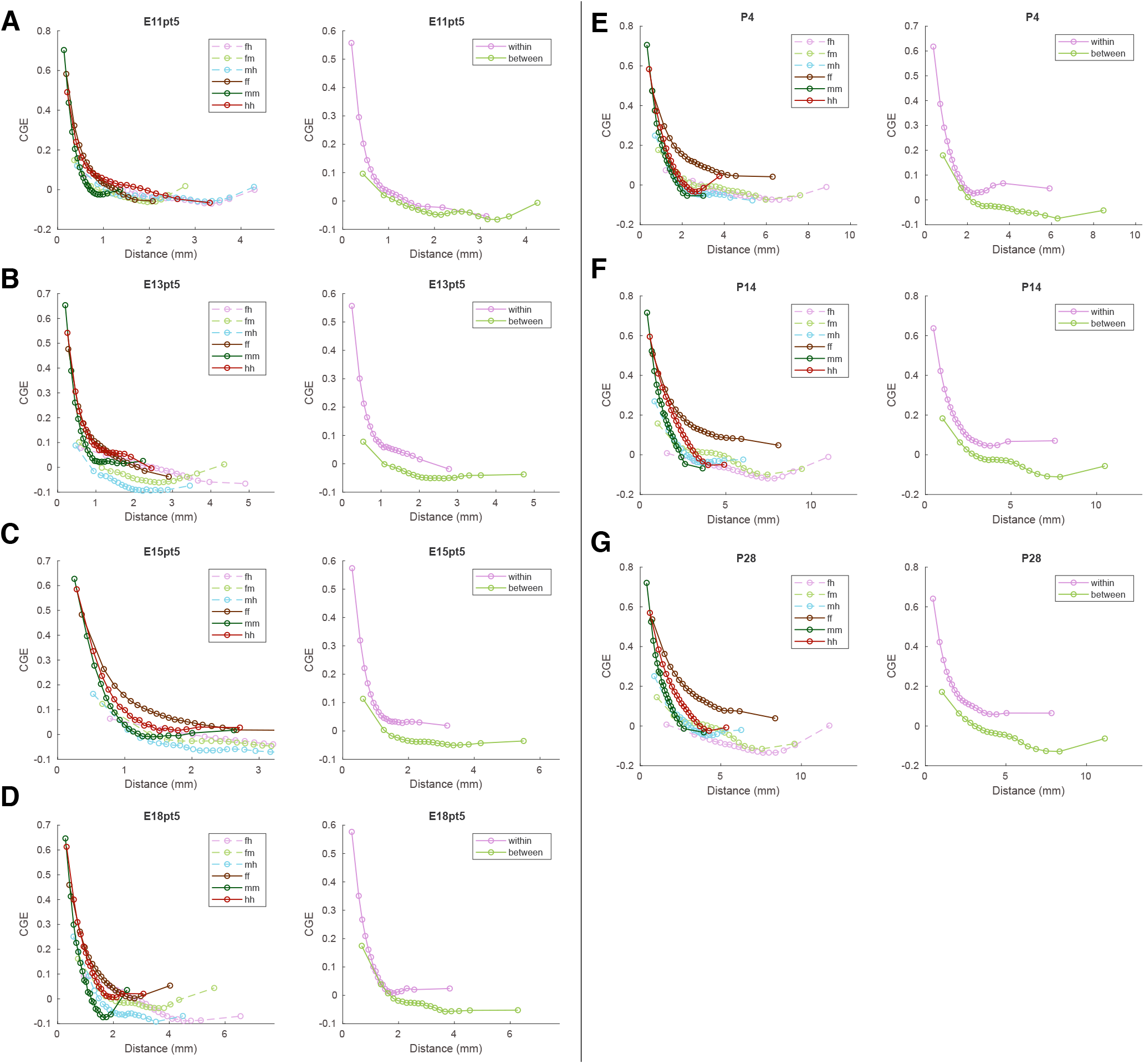
The distance-dependence of correlated gene expression (CGE) varies with broad anatomical divisions. We show the CGE(*d*) curves for each time point, as labeled: **A** E11.5, **B** E13.5, **C** E15.5, **D** E18.5, **E** P4, **F** P14, and **G** P28. In each panel, the left plot shows the CGE(*d*) across 20 equiprobable distance bins for a subset of at most 1000 voxels in each subdivision: forebrain (‘f’), midbrain (‘m’), and hindbrain (‘h’). CGE was computed for each type of voxel pair, the *within-division*: forebrain–forebrain (‘ff’), midbrain–midbrain (‘mm’), hindbrain-hindbrain (‘hh’); and *between–division*: forebrain-hindbrain (‘fh’), forebrain-midbrain (‘fm’), and midbrain–hindbrain (‘mh’). In the right panel of each subplot, within versus between division voxel pairs are agglomerated.

**Fig. S3.**
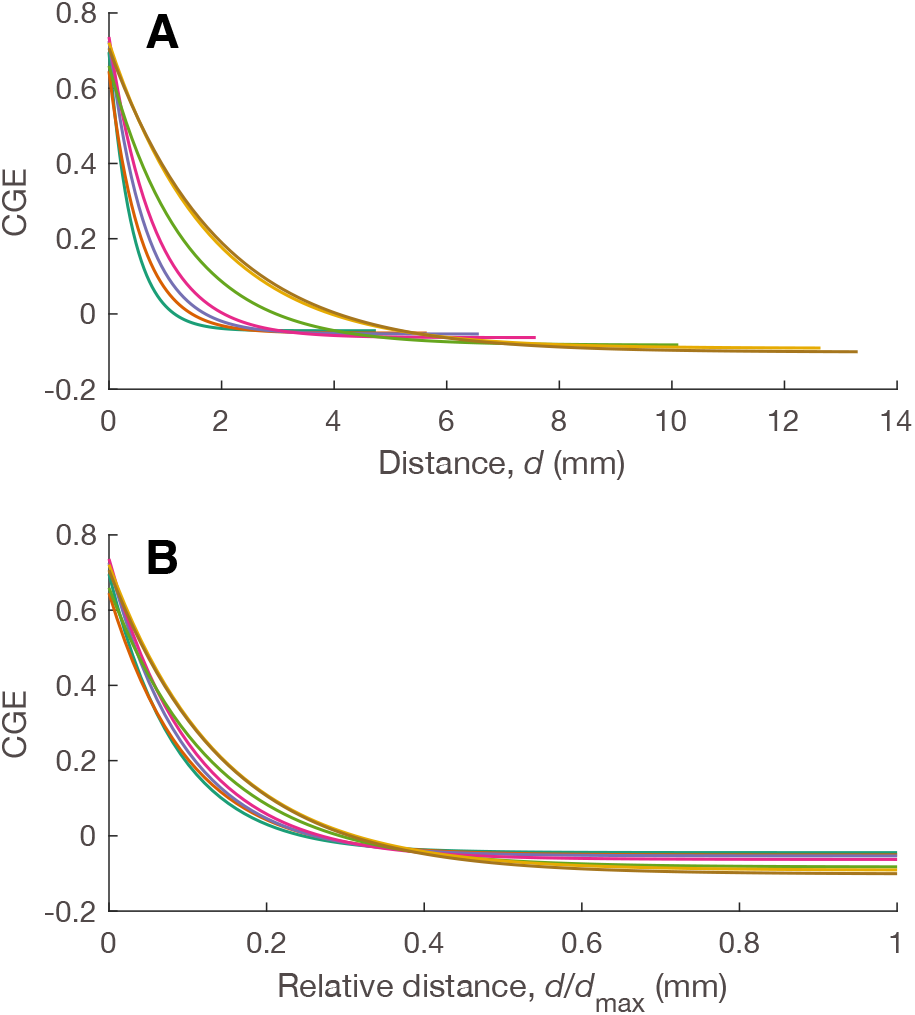
Rescaling distances by a linear measure of brain size, *d*_max_, reveals a similarity in the spatial embedding of correlated gene expression (CGE). Fitted exponential curves are plotted for all seven time points (colored as in Fig. 3) for: **A** Distance, *d*(mm), and **B** Relative distance, *d/d*_max_. The measure of brain size, *d*_max_, is taken as the maximum extent along the anterior–posterior axis.

**Fig. S4.**
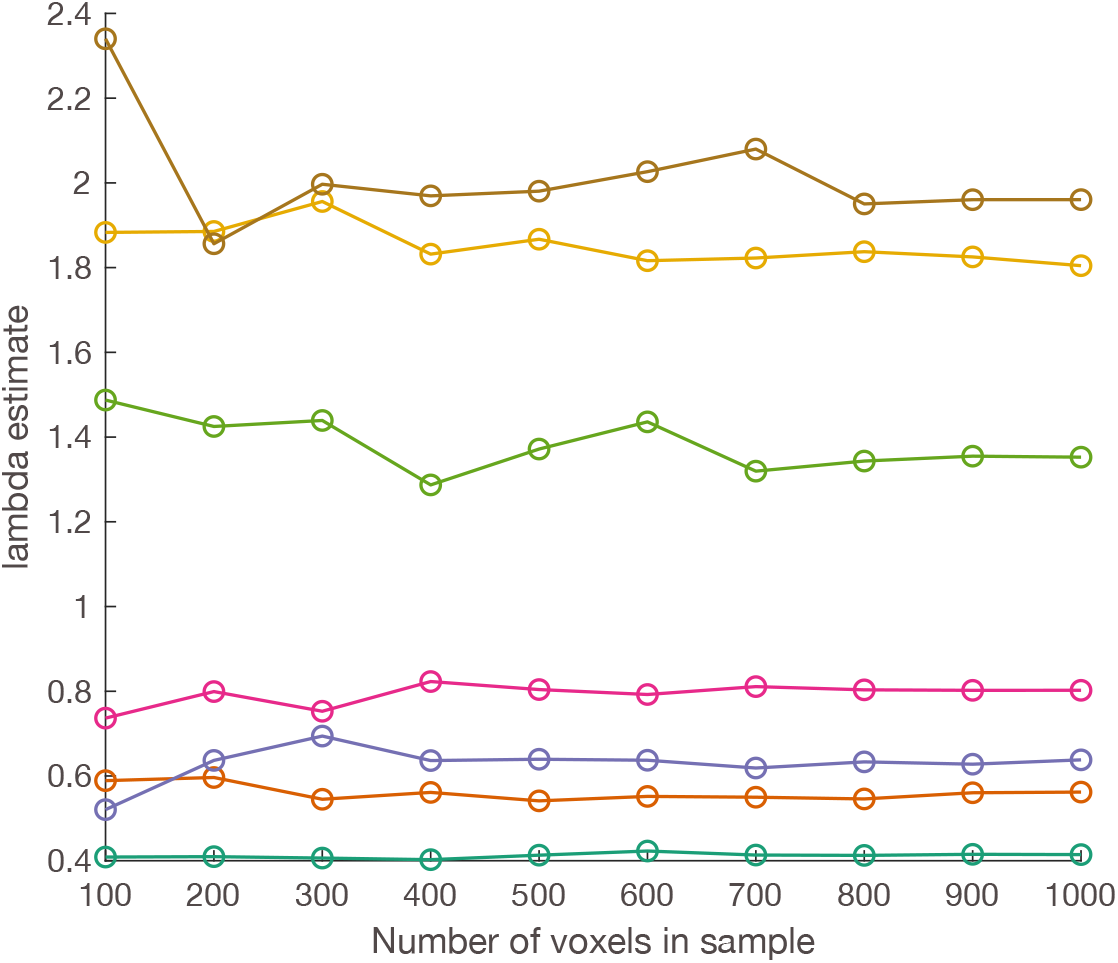
λ estimates are stable at random samples of 1000 voxels. We sampled different numbers of voxels (100 to 1000 in increments of 100), to estimate the spatial correlation length, λ, from an exponential fit.. Curves of the estimated λ values are plotted for each developmental time point using colors from the main text (see Fig. 3). The λ estimates across random voxel samples plateau around 500 voxels, and are stable at the 1000 voxel samples used here, indicating that 1000 is a sufficiently large sample size to obtain reliable λ estimates.

